# Social Housing Status Impacts Rhesus Monkeys’ Affective Responding in Classic Threat Processing Tasks

**DOI:** 10.1101/2021.05.16.444352

**Authors:** Joey A. Charbonneau, David G. Amaral, Eliza Bliss-Moreau

**Affiliations:** Neuroscience Graduate Program, University of California Davis; California National Primate Research Center, University of California Davis; The MIND Institute, University of California Davis School of Medicine; Department of Psychiatry and Behavioral Sciences, University of California Davis School of Medicine; Department of Psychology, University of California Davis

## Abstract

The established literature clearly demonstrates that whether or not monkeys are socially reared has long term consequences for their affective behavior. Yet, in the context of behavioral neuroscience and pharmacological studies, social context of adult animals is often ignored. When social context has been studied in adult monkeys, such studies have typically focused on welfare-related issues, as social isolation often leads to the development of abnormal behavior, rather than the impact on outcomes in behavioral neuroscience studies. Variation in social housing conditions for adult animals could have an impact on affective responding and may have significant implications for the interpretation of data from biopsychiatry and behavioral neuroscience studies. We evaluated the affective reactivity of rhesus monkeys (*Macaca mulatta*) maintained in one of four housing conditions (individually-housed, grate-paired, intermittently-paired, and continuously-paired) using two classic threat processing tasks—a test of responsivity to objects and the Human Intruder Test. Individually-housed monkeys exhibited consistently blunted sensitivity to ostensibly threatening stimuli as compared to socially-housed monkeys. Within the three socially-housed conditions, intermittently- and continuously-paired monkeys behaved similarly to each other and grate-paired monkeys exhibited relatively enhanced sensitivity to threatening stimuli. These findings suggest that the adult housing conditions of monkeys can robustly modulate affective responding in a way that may be consistent with behavioral phenotypes observed in human psychiatric conditions. Results are considered in the context of the broad behavioral and psychiatric neuroscience literatures, which have historically used individually-housed animals, pointing to the potential need to reconsider inferences drawn from those studies.

## Introduction

Nonhuman animals have been used for decades in the service of understanding the biological mechanisms of phenomena like mood and anxiety disorders and the basis of psychosocial behavior. Such studies often include neurobiological manipulations like the lesioning of particular brain regions or neural connections (e.g., [1]) or the pharmacological blockade or activation of particular receptors (e.g., [2]). Typically, such experiments rely on a comparison between manipulated and unmanipulated animals (“control animals”), and the behavior or task performance of control animals is assumed to be “healthy” or species “normative”. Such approaches typically ignore, or treat as inconsequential, aspects of the contexts in which animals are housed and tested, such as subjects’ socialization history and their social housing conditions, which potentially impact behavior. Contextual effects on control animals could influence how the effects of manipulations are interpreted and the underlying source of those effects may also interact with biological experimental manipulations. This complicates the interpretation of these data and the inferences drawn from ostensibly causal manipulations of biological mechanisms.

Neurobiological psychiatric research conducted with animals has typically focused on psychosocial outcomes (e.g., anxiety-like behavior, [3]) and measurements of specific behaviors thought to reflect these things (e.g., individual behaviors like freezing and cooing, [4]). Such behavioral measures can be sensitive to contextual effects (for reviews: [5,6]), sometimes in ways that are not immediately clear to experimenters. For example, the mere sex of the experimenter who was present resulted in significant differences in responses to a painful stimulus in both mice and rats such that responses to pain were inhibited in the presence of males [7]. This pain inhibition effect persisted across several different types of painful stimuli and was even found in the presence of just male-worn clothing [7].

The social contexts in which animals are reared and housed are increasingly recognized as playing an important role in psychosocial processing, just as social relationships and context are documented to play critical roles in human experience ([8]; for reviews: [9,10]). In adult humans, social isolation and loneliness are associated with increased mortality [11,12], chronic illness [13], and depression [14–16]. Beyond physical health, in humans, social context and the strength of relationships with family, neighbors, and community correlated with evaluations of subjective wellbeing [17].

The interplay between human social relationships and psychosocial processes likely has deep evolutionary roots, clearly seen in data gleaned with nonhuman primates, generally, and macaque monkeys, specifically. In macaques, social rearing conditions interact significantly with genotypic features to produce behavioral differences [18,19], to modify the density of serotonin receptors in the brain [20], and to influence hormonal responses to threat [21], among other observed differences (for a review, see [22]). Much of the literature on the effect of social context in monkeys has focused on the context in which infants are raised (see [23] for an early review of studies inducing depression in infant monkeys through social isolation and other means), despite increasing evidence that variation in social housing impacts adult monkeys as well [24,25]. Notably, in the vast majority of cases, adult monkeys who are housed alone (without direct contact with a social partner) are housed in rooms with other monkeys they can see and hear—so isolation in this case refers to physical features of social relationships. This visual, auditory, and olfactory contact is often used to qualify isolated housing on the basis that it is not truly isolated (e.g., [26,27]; for a discussion of this see [28]). Compared to socially-housed adult monkeys, however, individually-housed monkeys generate more aberrant behaviors, including more frequent self-injurious behavior [29,30], anxiety-like behaviors [3], and depression-like behaviors [31,32].

When removed from robust social contexts, the impact of social isolation appears quickly. When a cohort of adult male rhesus monkeys were relocated from semi-naturalistic, large outdoor social groups to individual indoor cages 69% (eighteen of twenty-six monkeys) exhibited depression-like behaviors in the first week after relocation [33]. The presence of a social partner upon relocation prevented an increase in the duration of such behaviors over time [34]. Despite evidence like this, social context effects have frequently been ignored in studies of adult monkeys, their behaviors, their brain activity, and their responses to manipulations thought to address psychosocial health. When the effects of individual housing have been studied beyond infancy, such studies have typically been motivated by the desire for better welfare (e.g., [35,36]) rather than to understand the potential mediating effects on other manipulations.

To assess the potential role of social context effects on behaviors commonly used to characterize changes in psychosocial health (including features related to depression and anxiety and other neurodevelopmental disorders), we analyzed responses of four cohorts of healthy adult rhesus macaques housed in various social configurations. Monkeys completed a test of responsivity to ostensibly threatening objects (as in [37–39]) and the Human Intruder Test (as in [40–42]). Here, we capitalize upon methodological consistency across cohorts (including standardized protocols for observation, interrater reliability, etc.), but variation in social housing conditions: individually-housed, grate-paired (able to access a partner partially through a metal grate), intermittently-paired (able to access a partner 7-8 hours/day), or continuously-paired (able to access a partner 24/7). Given the findings in the human mental health literature and nonhuman animal welfare literature, we expected that social housing condition would impact animals’ behavior during these tasks.

## Methods and Materials

Experimental procedures were approved by the University of California, Davis IACUC and were conducted in accordance with the National Institutes of Health guidelines for the use of animals in research. All procedures were carried out at the California National Primate Research Center (CNPRC).

### Subjects and Living Arrangements

Subjects were 22 adult male rhesus monkeys (*Macaca mulatta;* 5-10 years; Mean ± SD = 7.60 ± 1.37 years) who served as neurologically intact controls for a series of behavioral neuroscience experiments over the course of 10 years (2001-2011) in the same laboratory [39,43–48]. They were born at CNPRC and raised by their mothers in large social groups, remaining there through adulthood. They were relocated to temperature-controlled indoor rooms (12-hour light/dark cycle) where they were fed monkey chow (Ralston Purina, St. Louis, MO) twice daily supplemented with fresh fruit and vegetables twice per week and had *ad libitum* access to water and enrichment. Monkeys were housed in standard nonhuman primate caging (61 cm width x 66 cm depth x 81 cm height) in one of four possible configurations (see Table 1). Some data were previously published (A1-6: [48]; D1-7: [39]). All monkeys in the present report were relocated to indoor housing at least 3 months (Mean ± SD = 19.14 ± 4.58 months) prior to completing the tasks described here.

**Table 1:** Description of subjects

Housing configuration and variant used in ORT and HIT by monkey.
| Monkey | Housing Configuration | ORT | ORT Variant | HIT | HIT Variant |
| --- | --- | --- | --- | --- | --- |
| A1<sup>a,b,c,d</sup> | Individual | ✓ | 1 | ✓ | 1 |
| A2<sup>a,b,c,d</sup> | Individual | ✓ | 1 | ✓ | 1 |
| A3<sup>a,b,c,d</sup> | Individual | ✓ | 1 | ✓ | 1 |
| A4<sup>a,b,c,d</sup> | Individual | ✓ | 1 | ✓ | 1 |
| A5<sup>a,b,c,d</sup> | Individual | ✓ | 1 | ✓ | 1 |
| A6<sup>a,b,c,d</sup> | Individual | ✓ | 1 | ✓ | 1 |
| B1<sup>e</sup> | Grate | ✓ | 2 |  |  |
| B2<sup>e</sup> | Intermittent | ✓ | 2 |  |  |
| B3<sup>e</sup> | Intermittent | ✓ | 2 |  |  |
| B4<sup>e</sup> | Continuous | ✓ | 2 |  |  |
| C1<sup>c,f</sup> | Grate | ✓ | 3 | ✓ | 2 |
| C2<sup>c,f</sup> | Grate | ✓ | 3 | ✓ | 2 |
| C3<sup>c,f</sup> | Grate | ✓ | 3 | ✓ | 2 |
| C4<sup>c,f</sup> | Grate | ✓ | 3 | ✓ | 2 |
| C5<sup>c,f</sup> | Intermittent | ✓ | 3 | ✓ | 2 |
| D1<sup>g</sup> | Intermittent | ✓ | 4 | ✓ | 2 |
| D2<sup>g</sup> | Intermittent | ✓ | 4 | ✓ | 2 |
| D3<sup>g</sup> | Intermittent | ✓ | 4 | ✓ | 2 |
| D4<sup>g</sup> | Continuous (HIT),<br>Intermittent (ORT) | ✓ | 4 | ✓ | 2 |
| D5<sup>g</sup> | Continuous | ✓ | 4 | ✓ | 2 |
| D6<sup>g</sup> | Continuous | ✓ | 4 | ✓ | 2 |
| D7<sup>g</sup> | Continuous | ✓ | 4 | ✓ | 2 |

<sup>a</sup>[43]; <sup>b</sup>[44]; <sup>c</sup>[45]; <sup>d</sup>[48]; <sup>e</sup>[46]; <sup>f</sup>[47]; <sup>g</sup>[39]

### Behavioral Experiments

#### Object Responsivity Test

All twenty-two monkeys participated in one of four variants of the Object Responsivity Test (ORT; see Table 1). Only very similar or identical trials were compared and only dependent variables that were identical across experiments were analyzed. The order of testing was counterbalanced across test day for each cohort and all testing occurred within the same time frame relative to meals (i.e., monkeys tested across variants should have been equally hungry). All monkeys were tested in the same adapted lab care cage (80.01 cm x 83.32 cm x 101.6 cm) with a plexiglass front. The front had two vertical openings (25 cm x 5 cm) separated by 5.08 cm and centered 32.25 cm from the sides of the cage which allowed the monkeys to interact with objects and retrieve the food reward. An opaque guillotine door lowered by a rope-and-pulley system blocked the plexiglass between trials. All trials were 30 seconds.

Objects of varying complexity were used to evaluate changes in monkeys’ responses with variation in the affective value of stimuli (with simple objects being ostensibly less threatening than their complex counterparts; see discussion in [37]); complexity was varied across trials (Table 2). All monkeys completed complex object trials and so the responses on these trials were compared. Groups A, B, and D were exposed to both simple and complex object trials, so the responses on these trials were separately compared.

**Table 2.**
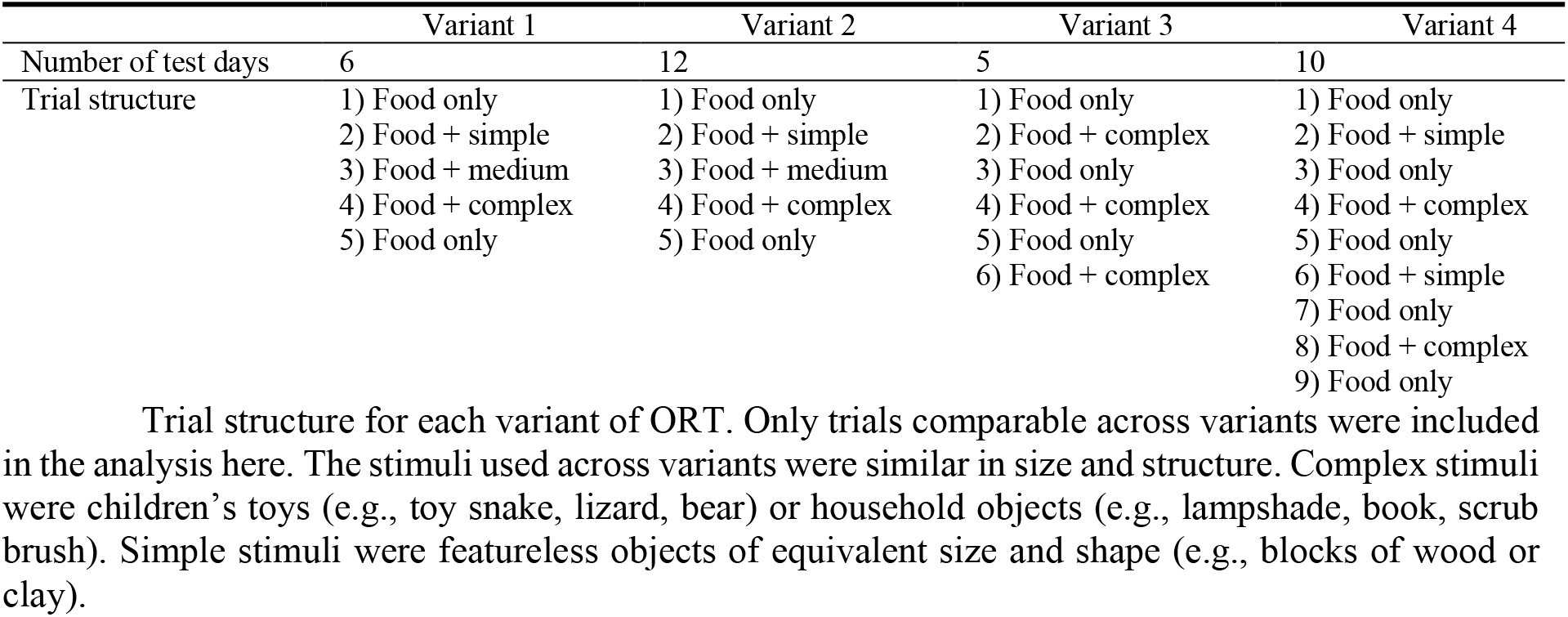
ORT trial structure by variant

Behavioral data collection was carried out by trained observers using The Observer (Noldus, Sterling, VA, USA) and included: 1) latency to retrieve the food reward; 2) frequency of food retrieval; 3) frequency of species-typical behaviors (lipsmack and grimace; summed to represent affective reactivity). Additional behaviors were recorded across each test administration but were not present in all data sets. All observers were trained to the lab standard of inter-rater reliability greater than 85%.

#### Human Intruder Test

Eighteen monkeys participated in one of two variants of the Human Intruder Test (HIT; see Table 1). Monkeys were tested individually in a standard primate cage (60 cm x 75 cm x 75 cm) and were exposed to an unfamiliar experimenter in four different conditions after a one-minute acclimation period as described previously by our group (e.g., [39,40,48,49]) and others (e.g., [41]). The four conditions were presented in the following order: 1) *profile far*: person facing 90 degrees away from the cage at 1 m distance; 2) *profile near*: profile at 0.3 m; 3) *stare far*: direct stare at the animal at 1 m; 4) *stare near*: stare at 0.3 m. Trials were 30 s (Variant 1) and 60 s (Variant 2). Tests were repeated across 5 days.

Behavioral data were scored live using a 1/0 sampling method by trained observers [as in [40,50]]. In Variant 1, six 5 second bins were used across 30 second trials. In Variant 2, six 10 second bins were used across 60 second trials. Behaviors that occurred within a bin received a score of 1 and behaviors that did not occur received a 0. To standardize across variants for data analysis, only the first three bins were used for monkeys in Variant 2 (totaling 30 seconds). Position in cage and six affect-related behaviors (i.e., lipsmack, grimace, threat, cage shake, tooth grind, and yawn) were scored. Affective behaviors were summed within and across bins to generate an index of affective reactivity.

### Statistical Analysis

R version 4.0.4 (R Core Team, 2020) was used for statistical analyses. Data were analyzed using linear mixed-effects (LME) models in *lme4* [51]. Follow-up LME models were fit to determine group differences in particular test conditions. Pairwise comparisons were made using estimated marginal means in *emmeans* [52]. Latency data were analyzed using a survival analysis (mixed effects Cox model implemented in *coxme* [53]) with a censored latency of 30 s. Difference scores were computed and analyzed for each animal using the HIT *stare near* and *stare far* data to investigate group differences across conditions. Analyses are relatively insensitive to unequal sample sizes [54]. Main analyses compared individually-housed monkeys against all socially-housed monkeys. Additional analyses compared the three socially-housed groups, presented below and in the supplemental materials.

## Results

### Experiment 1: Object Responsivity Test

#### Food retrieval

Retrieval frequencies during complex object trials were analyzed first because all subjects completed these trials. Individually-housed monkeys retrieved food significantly more frequently than socially-housed monkeys (*χ*^*2*^(1)=9.00, *p*<0.003); Figure 1A. Next, retrieval frequencies during simple and complex object trials were analyzed for the seventeen monkeys who completed both trial types, revealing the same effect of housing condition (*χ*^*2*^(1)=6.04, *p*<0.02). As expected, there was also a significant effect of object complexity (*χ*^*2*^(1)=11.11, *p*<0.001); all monkeys retrieved food more frequently during simple vs. complex object trials. The interaction of housing condition and complexity was not significant (*χ*^*2*^(1)=0.51, *p*=0.47); Figure 1B. There was a significant effect of housing condition on retrieval frequency (*χ*^*2*^(1)=7.01, *p*<0.04) when monkeys from the three social conditions were compared. Grate-paired monkeys retrieved less frequently than continuously-(*p*<0.04) or intermittently-paired (*p*=0.09) monkeys, which did not differ significantly from each other (*p*=0.99); Figure S1 and Supplemental Results.

**Figure 1.**
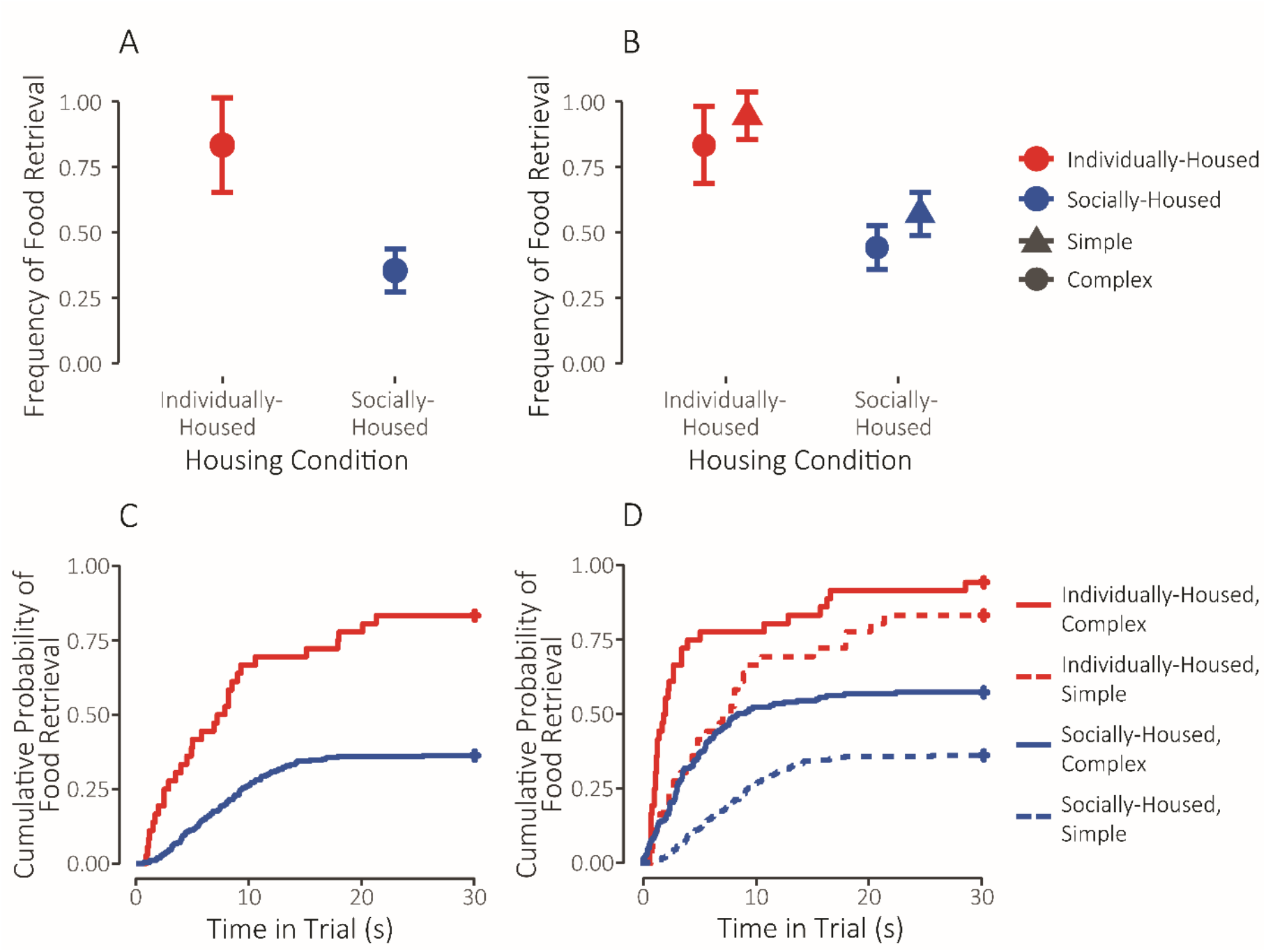
**(A)** Retrieval frequency during complex object trials for individually-(red, N=6) and socially-housed (blue, N=16) monkeys. **(B)** Retrieval frequency during simple (circle) and complex (triangle) object trials for individually-(red, N=6) and socially-housed (blue, N=11) housed monkeys. Means ± adjusted 95% confidence intervals are shown. **(C)** Cumulative probability of food reward retrieval as a function of time elapsed in trial during complex object trials for individually-(red, N=6) and socially-housed (blue, N=16) monkeys. **(D)** Cumulative retrieval probability during simple (dashed) and complex (solid) object trials for individually-(red, N=6) and socially-housed (blue, N=11) monkeys. Survival curves are shown. Retrieval latencies of 30 seconds (no retrieval) are right-censored.

Housing condition also influenced food retrieval latency. Individually-housed monkeys retrieved food significantly faster than socially-housed monkeys (*χ*^*2*^(1)=8.80, *p*<0.004) during complex object trials; Figure 1C. When simple and complex object trials were analyzed, there were significant main effects of housing condition (*χ*^*2*^(1)=7.50, *p*<0.007) and object complexity (*χ*^*2*^(1)=44.40, *p*<0.001) but not a significant interaction (*χ*^*2*^(1)=0.35, *p*=0.55). Individually-housed monkeys retrieved food faster than socially-housed monkeys in the presence of both object types. Both groups retrieved the food faster during simple vs. complex object trials; Figure 1D. There was a significant effect of housing condition on retrieval latency during complex object trials when monkeys from the three social conditions were compared (*χ*^*2*^(2)=7.11, *p*=0.03). Grate-paired monkeys retrieved food with longer latencies than either continuously-(*p*=0.02) or intermittently-paired (*p*=0.01) monkeys, which did not differ from each other (*p*=0.97). Figure S2 and Supplemental Results.

#### Affective reactivity

There was a significant effect of housing condition on affective reactivity to complex objects (*χ*^*2*^(1)=10.93, *p*<0.001). Individually-housed monkeys were less reactive to complex objects than socially-housed monkeys; Figure 2A. When responses to simple and complex objects were analyzed, the main effects of housing condition (*χ*^*2*^(1)=7.89, *p*<0.005) and complexity (*χ*^*2*^(1)=15.54, *p*<0.001) were both significant but the interaction was not (*χ*^*2*^(1)=0.05, *p*=0.82). Individually-housed monkeys were less reactive than socially-housed monkeys during both trial types. Both groups showed greater reactivity to complex vs. simple objects; Figure 2B. There was no significant effect of social housing condition on reactivity to complex objects when monkeys from the three social conditions were compared (*χ*^*2*^(2)=2.88, *p*=0.24); Figure S3 and Supplemental Results.

**Figure 2.**
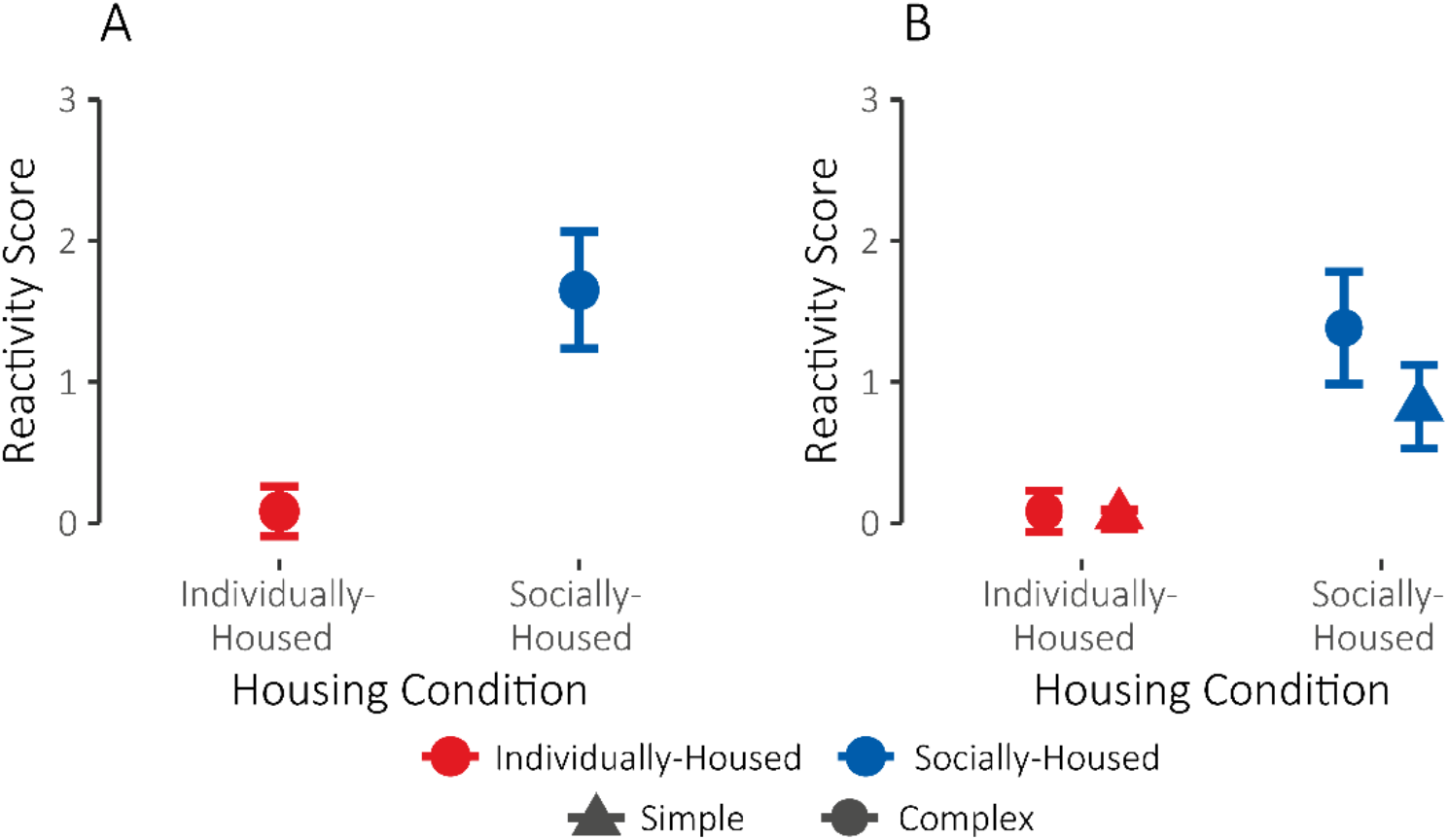
**(A)** Reactivity scores (combined frequency of affective behaviors) during complex object trials for individually-(red, N=6) and socially-housed (blue, N=16) monkeys. **(B)** Reactivity scores during simple (circle) and complex (triangle) object trials for individually-(red, N=6) and socially-housed (blue, N=11) monkeys. Means ± adjusted 95% confidence intervals are shown.

### Experiment 2: Human Intruder Test

#### Position in cage

There was a significant main effect of task condition on position at front of cage (*χ*^*2*^(3)=18.93, *p*<0.001) such that monkeys spent more time at the front of the cage during the *profile* vs. *stare* conditions. The main effects of test day (*χ*^*2*^(4)=1.47, *p*=0.83) and housing condition (*χ*^*2*^(1)=1.33, *p*=0.25) were not significant. There was a significant interaction of housing condition and task condition (*χ*^*2*^(3)=22.26, *p*<0.001). Comparison of the estimated marginal means revealed significant differences between groups only in the *stare near* condition (*p*<0.001; *profile far*: *p*=0.30; *profile near*: *p*=0.16; *stare far*: *p*=0.17). Individually-housed monkeys spent more time at the front of the cage during this condition than socially-housed monkeys; Figure 3A. There was no main effect of social housing condition on position in cage when monkeys from the three social conditions were compared (*χ*^*2*^(2)=2.02, *p*=0.36); Figure S4 and Supplemental Results.

**Figure 3.**
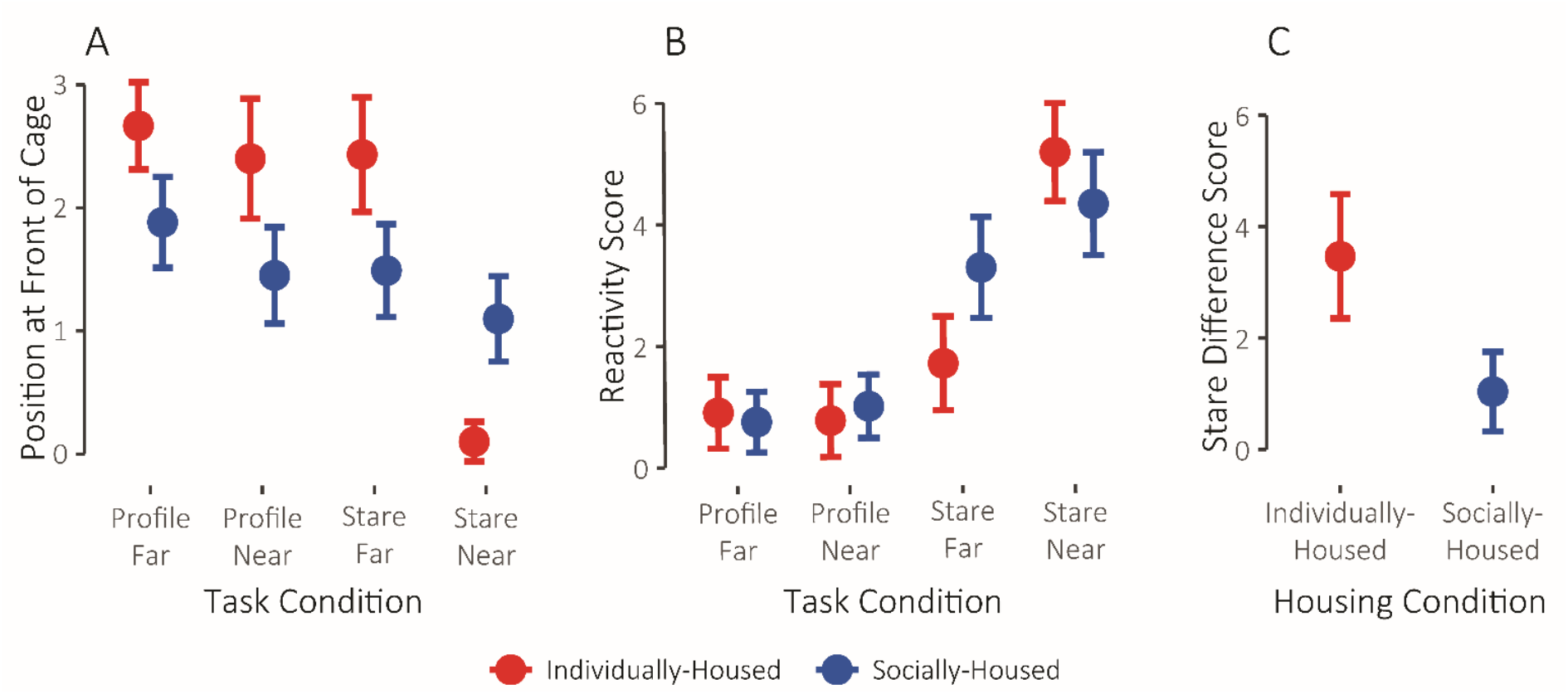
**(A)** Tendency to be present at the front of the cage during the four Human Intruder Test conditions for individually-(red, N=6) and socially-housed (blue, N=12) monkeys. **(B)** Reactivity scores (combined frequency of affective behaviors) during the four Human Intruder Test conditions for individually-(red, N=6) and socially-housed (blue, N=12) monkeys. **(C)** Difference scores (stare near – stare far) for individually-housed (red, N=6) and socially-housed (blue, N=12) monkeys. Means ± adjusted 95% confidence intervals are shown.

#### Affective reactivity

There were significant main effects of test day (*χ*^*2*^(4)=16.55, *p*<0.003) and task condition (*χ*^*2*^(3)=291.34, *p*<0.001) on affective reactivity. The main effect of housing condition was not significant (*χ*^*2*^(1)=0.17, *p*=0.68). However, the interaction between housing condition and task condition was significant (*χ*^*2*^(3)=20.98, *p*<0.001); Figure 3B and Figure S5.

To understand what was driving the significant interaction, we analyzed responses in the *profile* and *stare* conditions separately, as monkeys are often not reactive during *profile* [40]. As expected, reactivity was low during the *profile* conditions and there was not a significant effect of housing condition (*χ*^*2*^(1)=0.63, *p*=0.43), test day (*χ*^*2*^(4)=1.80, *p*=0.77) or task condition (*χ*^*2*^(1)=0.57, *p*=0.45), nor was the interaction between these significant (*χ*^*2*^(1)=1.63, *p*=0.20). When reactivity during *stare* trials was considered, there were significant main effects of test day (*χ*^*2*^(4)=14.74, *p*<0.006) and task condition (*χ*^*2*^(1)=36.00, *p*<0.001), but not housing condition (*χ*^*2*^(1)=0.15, *p*=0.70). Critically, the interaction between housing condition and task condition was significant (*χ*^*2*^(1)=19.34, *p*<0.001). Individually-housed monkeys were much more reactive in *stare near* than *stare far*, while socially-housed monkeys showed very similar reactivity across these two conditions. Comparison of the estimated marginal means did not reveal a significant difference between individually- and socially-housed monkeys in either *stare far* (*p*=0.36) or *stare near* (*p*=0.32). As the significant interaction between housing condition and task condition suggested that socially- and individually-housed monkeys calibrated their responses differently across conditions, we computed difference scores between *stare near* and *stare far* responses for each monkey. Analysis of the difference scores revealed a significant effect of housing condition (*χ*^*2*^(1)=22.30, *p*<0.001). The difference between conditions was greater for individually-as compared with socially-housed monkeys; Figure 3C. There was no main effect of social housing condition on reactivity when monkeys from the three social conditions were compared (*χ*^*2*^(2)=3.16, *p*=0.21); Figure S6 and Supplemental Results.

## Discussion

Nonhuman primates are used extensively in psychiatric neuroscience research [4,55,56], predicated on the importance of homologies both in psychosocial functions and neuroanatomy across monkeys and humans [57,58] and the desperate need for new treatments and interventions for psychiatric conditions as global incidence soars [59,60]. Despite many decades of use as in such experiments, the social context in which subject monkeys live and the large impact that social context may have on psychosocial behavior has historically gone largely unacknowledged; if acknowledged, it has typically been in terms of animal welfare and not experimental outcomes (e.g., [61,62]). Our data demonstrate that social context matters for the very outcomes about which behavioral neuroscientists, pharmacologists, and physiologists interested in psychiatric diseases care. Monkeys who were individually-housed during experiments generated very different affective responses to threat compared to monkeys who had some form of social contact. Patterns of behaviors were consistent across experiments relative to housing condition.

During ORT, individually-housed monkeys showed dampened responses to threat, retrieving food rewards from beside threatening objects faster, more frequently, and with less reactivity than socially-housed monkeys. These data are consistent with the ideas that a transition from social groups to individual housing induces a depressive phenotype in animals [33,34,63,64] and that depression in humans induces threat insensitivity [65]. Similarly, during HIT, individually-housed monkeys were both less reactive and more willing to spend time at the front of the cage than socially-housed monkeys in three of the four conditions. In the 4^th^ condition, *stare near*, individually-housed monkeys exhibited elevated threat-responding, suggesting they may have a different threshold for response than socially-housed monkeys. Importantly, there was variance in the responses of monkeys housed in different social housing contexts such that monkeys with restricted social access (grate pairing) behaved differently from monkeys who had full physical contact in that they were more reactive and retrieved food rewards less frequently and more slowly during ORT (but were comparable in HIT). It is important to note that the two tasks used here, ORT and HIT, are widely used to characterize affective reactivity of monkeys’ natural or induced phenotypes (e.g., [38,41,66–68]) and are thought to provide “ground truth” about an individual’s affective reactivity.

Our data make clear that social housing conditions during adulthood shape affective responding in a way that is similar to how early social housing conditions have a profound impact on affective responding. More than 50 years ago Harlow and colleagues found that social isolation in infancy disrupts normal patterns of social and exploratory behaviors [23,69,70] and that these disruptions persist beyond the period of isolation. While many studies of social restriction in infancy focus on the impact of rearing babies in nurseries without their mothers (typically with humans, or with other caretakers such as dogs [71]), accumulating evidence demonstrates that infants raised alone with their mothers or only with other mother-infant pairs have disrupted behavior and physiology compared to infants raised in social groups (e.g., increased heart rates and lower respiratory sinus arrhythmia in [72]; reductions in immune response in [73]). Several studies have shown that variation in behavior during HIT relates to social rearing conditions: peer-reared monkeys exhibited less exploratory and more anxiety-like behaviors than mother-reared monkeys [74]; both indoor mother-reared and nursery-reared monkeys exhibited greater affective reactivity than outdoor socially-reared monkeys [41]. Nursery-reared, relative to mother-reared, monkeys also differ neurobiologically and physiologically, including altered functioning of the noradrenergic system [75], changes to white matter structures like the corpus callosum [76], disruptions in the development of the serotonergic system [63], and altered functioning of the HPA axis [77]. We know that reducing social contact can generate a depressive phenotype in a significant portion of adult monkeys [33,34,78] although this has largely been treated as fleeting or recoverable. Those data, with the infant rearing condition data, and our data, suggest that social contexts in adulthood have a similar impact on psychosocial outcomes and have simply gone uninvestigated because of norms related to husbandry in behavioral neuroscience.

What is also clear from our data is that not all social housing provides the same social context. In particular, grate-pairing appears to be different from other social housing conditions. This parallels what has been demonstrated in the infant literature, where infants who are raised “socially” with other infants or their mothers alone, but not in groups, have different behavioral and neurophysiological phenotypes from each other and group reared infants (for a review, see [22]). The behavior of grate-paired monkeys differed monkeys who had full continuous or intermittent contact. For example, during complex object trials, grate-paired monkeys were most reactive and retrieved the food reward less frequently than other socially-housed monkeys. This was the opposite pattern of results generated by individually-housed monkeys. One hypothesis is that grate-pairing causes an anxiety-like phenotype while single-housing causes a depression-like phenotype, pulling animals in those conditions away from normal in opposite directions.

Many neuropsychiatric and behavioral neuroscience studies have been conducted with singly housed monkeys, including studies probing the neural basis of socioaffective processing [1,45,79–81] as well as the basis of addiction and drug-seeking behaviors [82–86]. Even studies intending to specifically address the ability to induce depression in monkeys though means other than social isolation (e.g., administration of the immune cytokine interferon-alpha [87]; exposure to social stress [78]) and those intending to determine the impact of early social deprivation [88] or stress [89], have all used monkeys that were singly housed as adults. In light of the findings we present here, the documented effects of these manipulations may be severely confounded by the impact of restricted social contact in adulthood, which has important implications for the interpretation of the results of these studies and the many others that have used individually-housed monkeys. Conclusions drawn from such studies must be reconsidered with regards to whether or not the behavior of control animals is normative. Taking the impact of social context seriously may provide a beneficial opportunity to resolve conflicts in the literature (e.g., the role of the amygdala in social cognition, as discussed by [90]).

Our analyses benefited from access to data from experiments that were carried out using very similar protocols and monkeys in the same laboratory. Given methodological consistencies across studies, it is unlikely that the effects are due solely to differences in the ways that the experiments were carried out. Importantly, our subjects were raised in conditions most closely resembling the natural rearing conditions of a macaque monkey as are possible in a captive setting through adulthood (i.e., ∼4 years of age; [91]). Subjects were chosen for these experiments because they were ‘normal’ and exhibited all of the expected species-typical behaviors when observed in the field cages. It is therefore striking that we observed as strong of an impact of adult social condition as we did. Nevertheless, future studies should replicate the present experiments in a group of monkeys housed in different social contexts (both between and within subjects) and tested concurrently by the same experimenters.

Causal manipulations in nonhuman animals’ brains are capable of providing mechanistic insights into the neural basis of human psychiatric diseases [56,92]. Ultimately, such studies are forced to draw inferences based on comparisons between control and manipulated animals and often assume that the behavior of controls is normal or healthy. If controls have behavioral phenotypes consistent with neuropsychiatric disorders like anxiety or depression, then these subjects do not provide a faithful baseline. Further, the conditions that drive controls to take on this altered phenotype may interact with experimental manipulations leading to incorrect inferences about the impact of the manipulation. While we demonstrate the impact of social context, other aspects including diet, circadian rhythms, quality of social relationships, husbandry practices, and features of experimenters may drive similar changes in behavior. The ultimate aim of this work is to initiate a conversation regarding the potential impact of context on the results of studies aiming to reveal the nature or pathophysiology of neuropsychiatric disorders and to rethink existing inferences in the literature. We hope that such a conversation, in tandem with additional experiments specifically testing contextual effects, will be integral to further investigation of mood disorders, their basis, and their treatment.

## Acknowledgements

The authors wish to thank Jeffrey Bennett for his help finding archival data records.

## Author Contributions

JAC, EBM, and DGA were responsible for the study concept and design. JAC and EBM assisted with data analysis and interpretation of findings. JAC drafted the manuscript. EBM and DGA provided critical revision of the manuscript for important intellectual content. All authors reviewed content and approved the final version for publication. All authors agree to be accountable for all aspects of the work to ensure that questions related to the accuracy or integrity of any part of the work are appropriately investigated and resolved.

## Supplemental Results

### Object Responsivity

All three groups of socially-housed monkeys (grate-paired, intermittently-paired, and continuously-paired) were compared on responses to complex object trials. The effects of social housing condition in these analyses are presented in the main text. The responses of intermittently- and continuously-paired monkeys were also compared for both simple and complex object trials. Grate-paired monkeys were not included in these comparisons as only one grate-paired monkey completed both of these trial types.

#### Food retrieval

There was not a main effect of object complexity (*χ*^*2*^(1)=3.54, *p*=0.06) or social housing condition (*χ*^*2*^(1) = 0.07, *p* = 0.79) on retrieval frequency when intermittently- and continuously-paired monkeys were compared on simple and complex object trials. The interaction between object complexity and housing condition was also not significant (*χ*^*2*^(1)=0.30, *p*=0.59). See Figure S1B. There was a main effect of object complexity (*χ*^*2*^(1)=23.88, *p*<0.001) but not social housing condition (*χ*^*2*^(1)<0.001, *p*=0.99) on latency to retrieve the food for these groups. The interaction between complexity and social housing condition was also not significant (*χ*^*2*^(1)=0.001, *p*=0.97). Both groups took longer to retrieve the food during complex vs. simple object trials. See Figure S2B.

#### Affective reactivity

There was no significant effect of social housing condition on reactivity to simple and complex objects for the intermittently- and continuously-paired groups (*χ*^*2*^(1)=0.07, *p*=0.80). However, as expected, the main effect of complexity was significant (*χ*^*2*^(1)=13.49, *p*<0.001). Across both conditions, monkeys were more reactive to complex objects as compared to simple ones. See Figure S3B.

### Human Intruder Test

The main effects of social housing condition on responses to the HIT are presented in the main text. The effects of test day and task condition are presented here. These comparisons include the grate-paired, intermittently-paired, and continuously-paired monkeys.

#### Position in cage

The main effect of test day on position in cage was not significant (*χ*^*2*^(4)=3.55, *p*=0.47). The effect of task condition was significant (*χ*^*2*^(3)=10.01, *p*=0.02). The interactions between task condition and test day (*χ*^*2*^(8)=3.48, *p*=0.90) or housing condition (*χ*^*2*^(6)=7.20, *p*=0.30) were also not significant. All three groups showed a decline in tendency to be present at the front of the cage as the task condition became more ostensibly threatening (i.e., they tended to be present at the front more during the *profile* conditions as compared to *stare* and within these more during *far* as compared to *near*). See Figure S4.

#### Affective reactivity

The main effects of test day (*χ*^*2*^(4)=15.00, *p*<0.005) and task condition (*χ*^*2*^(3)=154.36, *p*<0.001) on affective reactivity were significant. These effects followed the expected pattern with less reactivity across test days and more reactivity in the *stare* conditions as compared to *profile*. See Figure S6.

**Figure S1.**
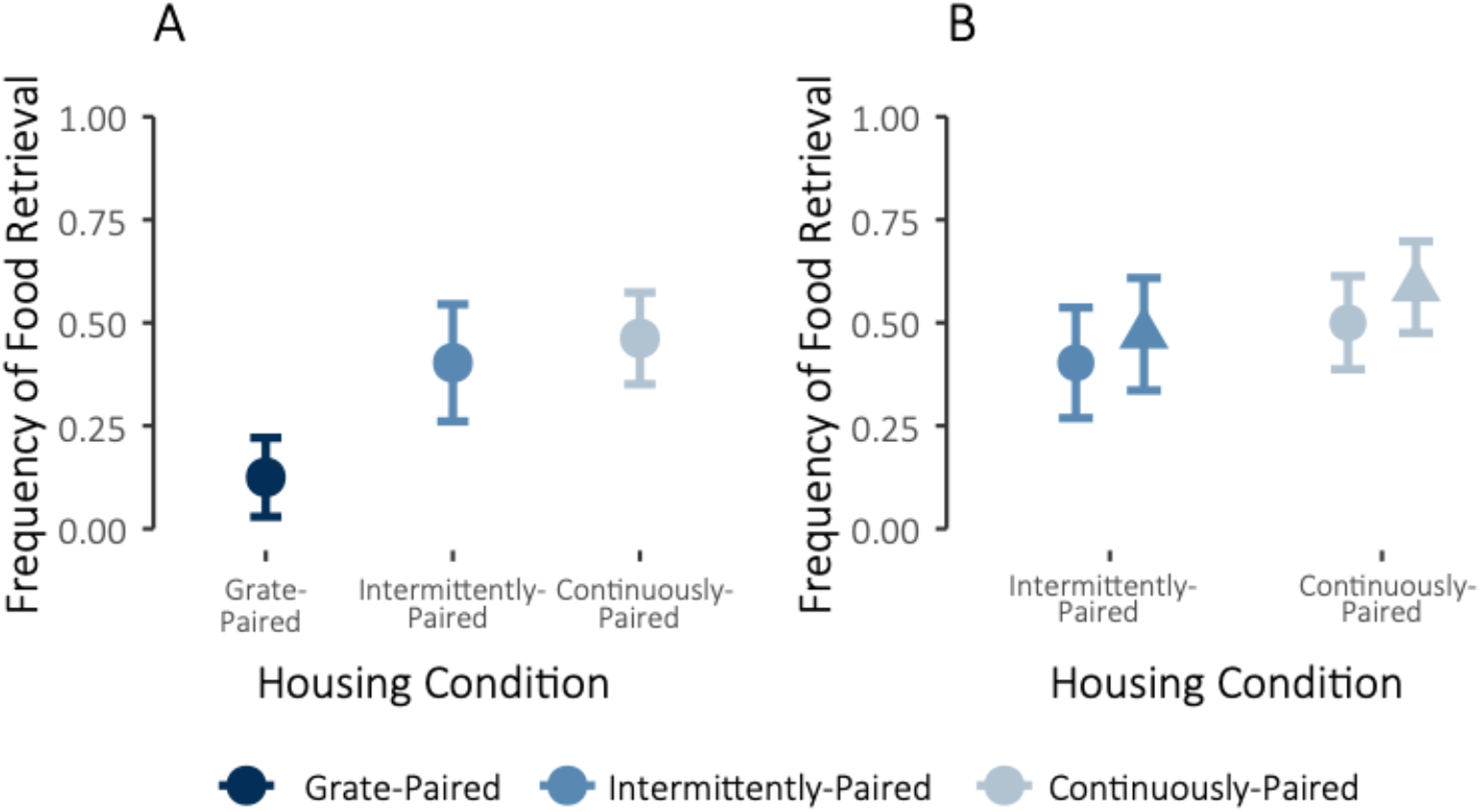
**(A)** Retrieval frequency during complex object trials for grate-paired (dark blue, N=5), intermittently-paired (medium blue, N=7), and continuously-paired (light blue, N=4) monkeys. **(B)** Retrieval frequency during simple (circle) and complex (triangle) object trials for intermittently-paired (medium blue, N=6) and continuously-paired (light blue, N=4) monkeys. Means ± adjusted 95% confidence intervals are shown.

**Figure S2.**
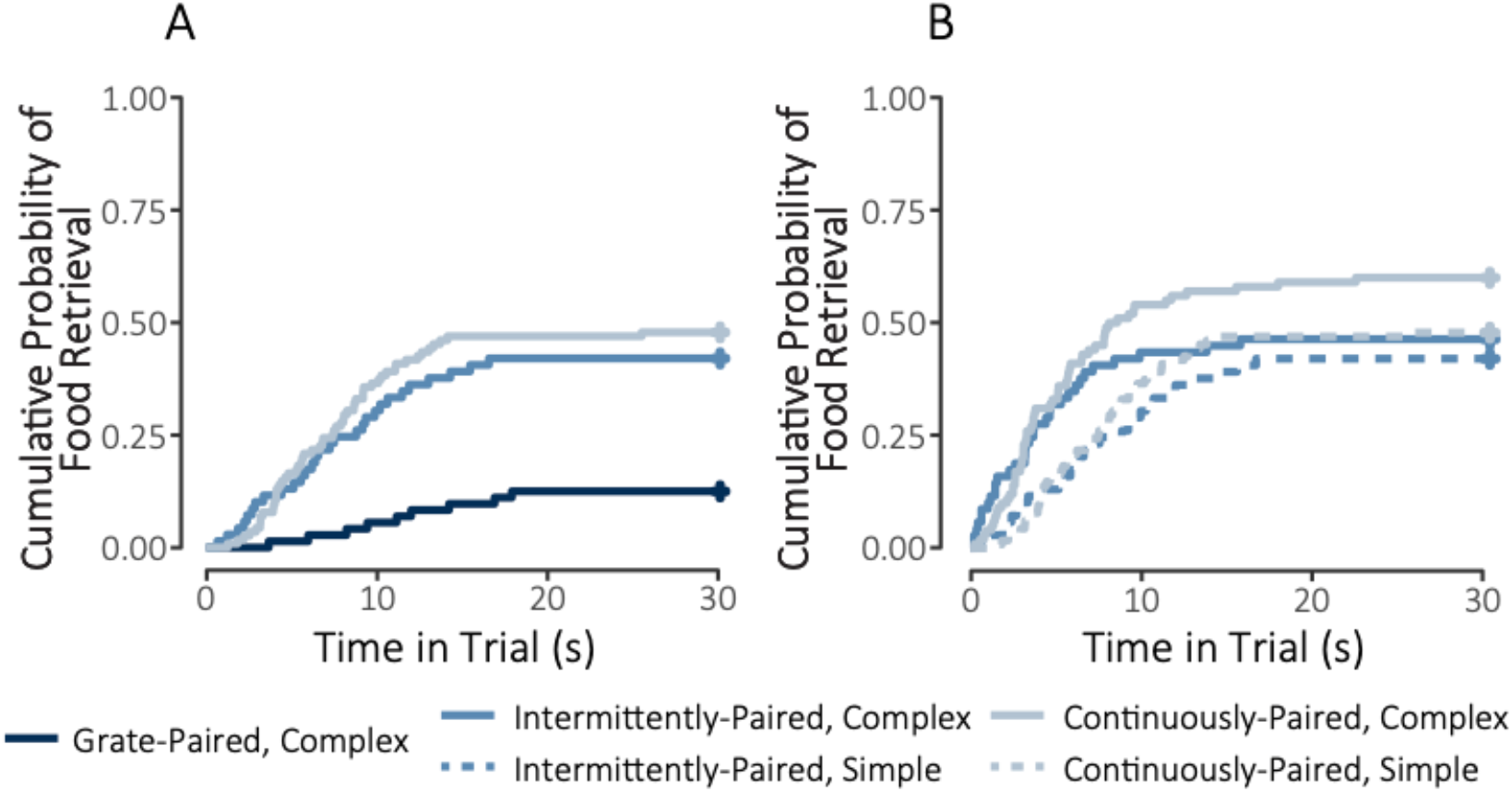
Cumulative probability of food reward retrieval as a function of time elapsed in trial. **(A)** Retrieval probability during complex object trials for grate-paired (dark blue, N=5), intermittently-paired (medium blue, N=7), and continuously-paired (light blue, N=4) monkeys. **(B)** Retrieval probability during simple (dashed) and complex (solid) object trials for intermittently-paired (medium blue, N=6) and continuously-paired (light blue, N=4) monkeys. Survival curves are shown. Retrieval latencies of 30 seconds (no retrieval) are right censored.

**Figure S3.**
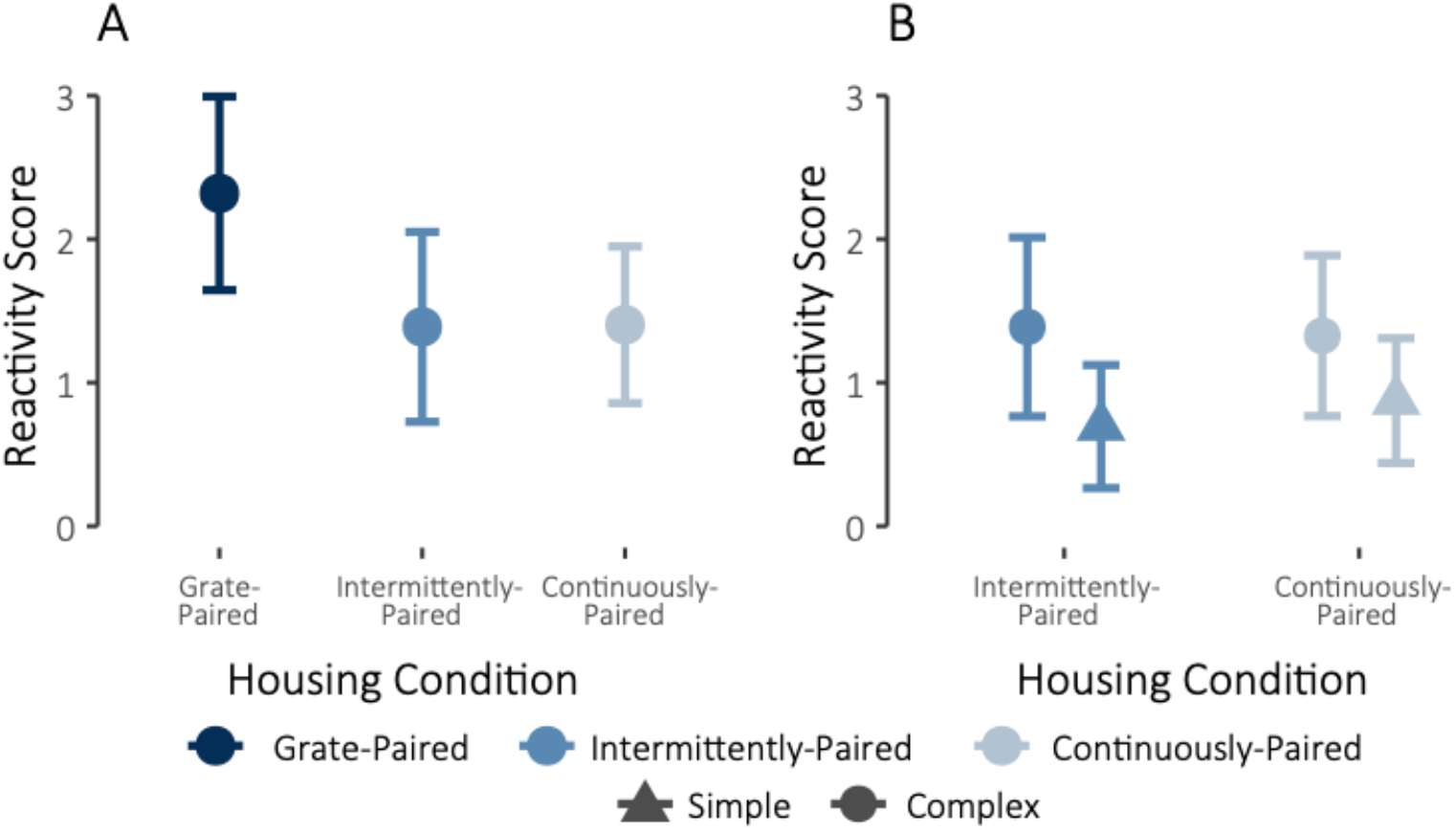
**(A)** Reactivity scores during complex object trials for grate-paired (dark blue, N=5), intermittently-paired (medium blue, N=7), and continuously-paired (light blue, N=4) monkeys. **(B)** Reactivity scores during simple (circle) and complex (triangle) object trials for intermittently-paired (medium blue, N=6) and continuously-paired (light blue, N=4) monkeys. Means ± adjusted 95% confidence intervals are shown.

**Figure S4.**
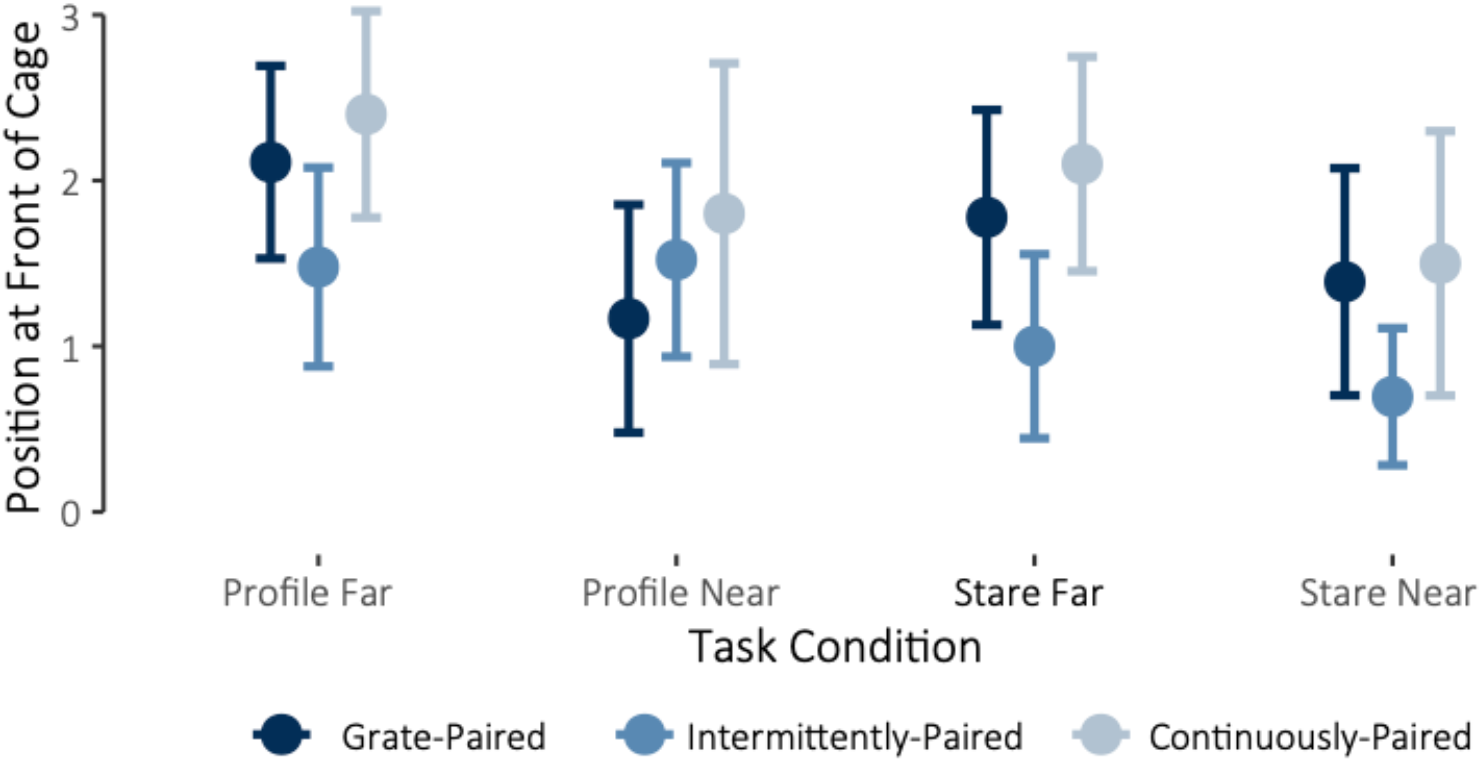
Tendency to be present at the front of the cage during the four Human Intruder Test conditions for grate-paired (dark blue, N=4), intermittently-paired (medium blue, N=7), and continuously-paired (light blue, N=4) monkeys. Means ± adjusted 95% confidence intervals are shown.

**Figure S5.**
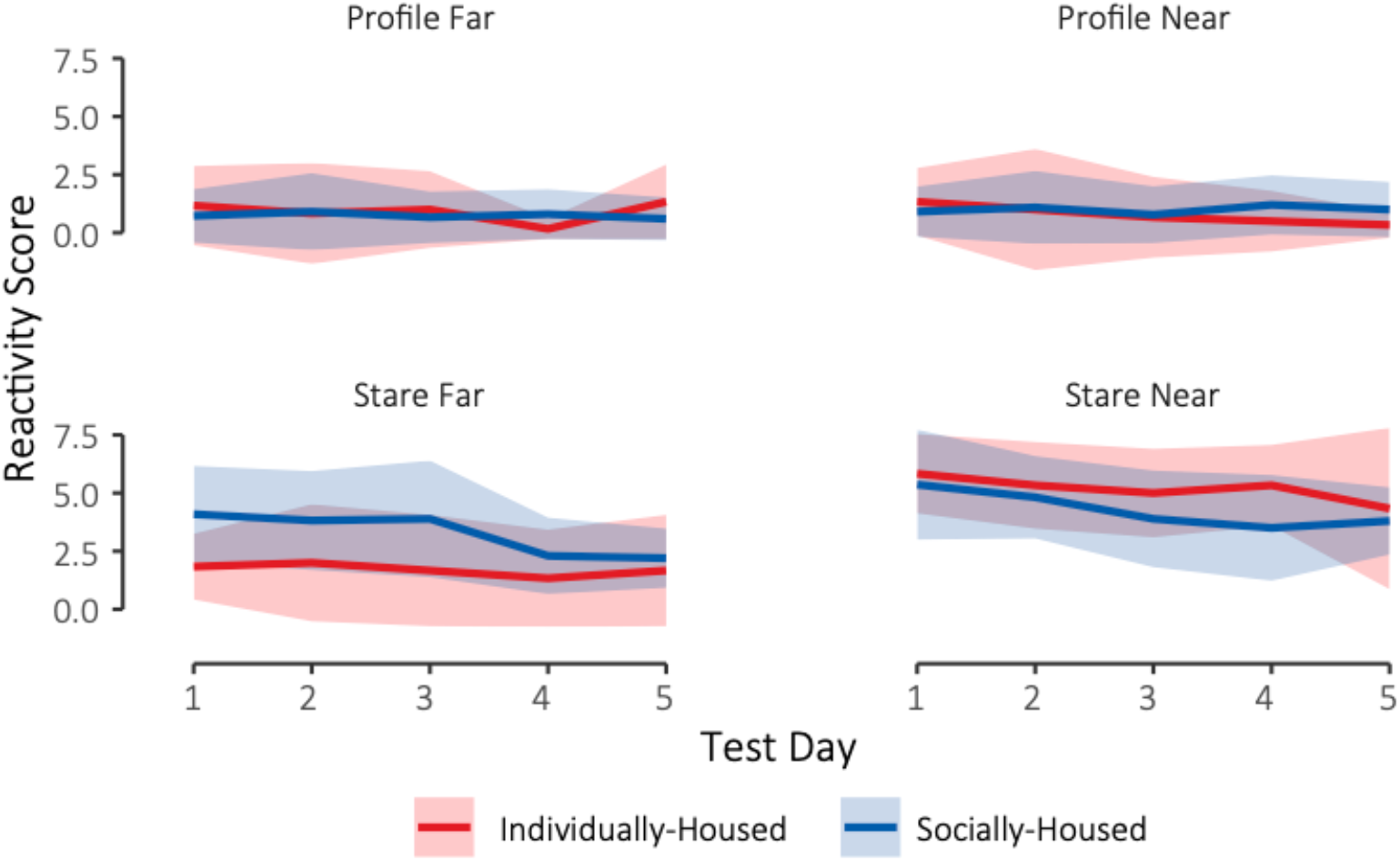
Reactivity scores during the four Human Intruder Test conditions for individually-housed (red, N=6) and socially-housed (blue, N=12) monkeys by test day. Means ± adjusted 95% confidence intervals are shown.

**Supplementary Figure 6.**
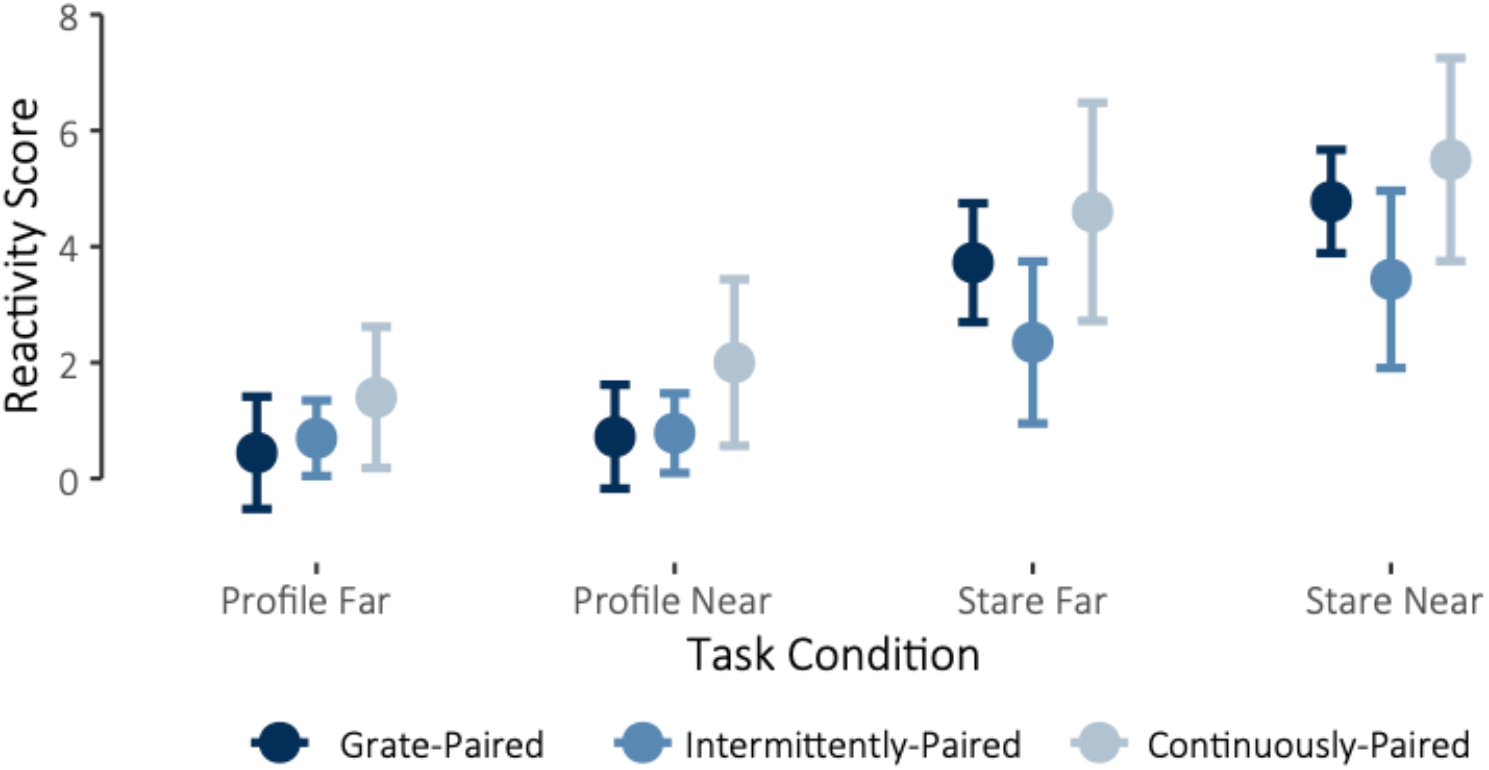
Reactivity scores during the four Human Intruder Test conditions for grate-paired (dark blue, N=4), intermittently-paired (medium blue, N=7), and continuously-paired (light blue, N=4) monkeys. Means ± adjusted 95% confidence intervals are shown.

